# Photoacoustic Fingerprinting for Robust Molecular Imaging

**DOI:** 10.64898/2026.04.13.718141

**Authors:** Colton McGarraugh, Luca Menozzi, Rui Yao, Dora Eng-Wu, Van Tu Nguyen, Soon-Woo Cho, Samuel Francis, Junjie Yao

## Abstract

Quantitative molecular imaging in photoacoustics is fundamentally limited by the ill-posed nature of spectral unmixing, where spectral overlap, noise, and unknown fluence introduce bias in conventional inversion-based methods. We introduce photoacoustic fingerprinting (PAF), a framework that reframes spectral unmixing as a fingerprint recognition problem. PAF interprets multispectral signals as high-dimensional fingerprints encoding both molecular composition and measurement distortions. Inspired by magnetic resonance fingerprinting, PAF uses a recurrent neural network trained on synthetic data spanning realistic mixtures, noise levels, and fluence variations to directly infer molecular concentrations from spectral shape. PAF enables accurate and robust quantification in regimes where conventional methods break down, including low signal-to-noise conditions, spectrally correlated mixtures, and unknown fluence distortions. In controlled simulations, PAF consistently outperformed non-negative least squares, with the largest gains observed for spectrally overlapping chromophores such as collagen. In phantom studies, PAF improved molecular specificity by correctly localizing collagen and recovering water contrast despite similar spectral reconstructions. In *ex vivo* mouse livers, PAF detected lipid accumulation associated with steatosis, and in human arteries, it identified molecular signatures consistent with thrombus and lipid-rich plaque. These results establish PAF as a generalizable framework for label-free molecular imaging and a promising step toward quantitative photoacoustic diagnostics.

## 1. Introduction

Photoacoustic imaging (PAI) has uniquely positioned itself at the intersection of rich optical contrast and high acoustic resolution, offering the potential to map deep-tissue molecular composition non-invasively. [1, 2]. By detecting ultrasound waves generated from pulsed optical absorption, PAI promises to quantify endogenous chromophores (hemoglobin, lipids, water, and collagen) beyond the diffusion limit of pure optical microscopy [1–3]. However, accurate molecular quantification in photoacoustic imaging remains fundamentally limited by the ill-posed nature of spectral unmixing, particularly in deep tissue settings where optical heterogeneity and measurement noise are unavoidable [2]. In practical imaging, wavelength-dependent fluence and noise are unknown, spatially heterogeneous, and often unmeasurable, rendering conventional inversion-based approaches intrinsically biased and often unreliable in practical imaging scenarios.

Despite challenges, an understanding of the absorption coefficient spectra of chromophores has enabled numerous photoacoustic applications, most notably the characterization of blood oxygen saturation [2, 4]. By imaging blood samples at multiple wavelengths, hemoglobin oxygen saturation (sO_2_) can be estimated through spectral unmixing [5]. Several *in vivo* studies have applied this principle to research tumor development through hypoxia [6], metastatic lymph node detection [7], and ischemic stroke [8]. The accuracy of these reconstructions strongly influences the reliability of such physiological measurements; therefore, the choice in unmixing method is important. Several approaches have been proposed for spectral unmixing: the linear model [2, 8, 9], the transport based model [2], the acoustic spectrum based model [10], and the absorption-saturation based model [4]. These methods rely on explicit modeling of light transport or linear inversion, which are either computationally prohibitive, limited in resolution, or fail to generalize under realistic imaging conditions.

Among these approaches, the linear model is the most used due to its simplicity and ease of implementation. However, this model assumes uniform optical fluence and linear relationships between optical absorption and measured signal intensity. In practice, these assumptions break down in deep tissues and regions of increased optical heterogeneity. The depth at which fluence non-uniformity becomes dominant depends on tissue optical properties and the imaging geometry. Therefore, fluence non-uniformity introduces significant bias in quantitative unmixing if not accounted for (e.g., iterative fluence-compensation methods) [2, 11, 12]. As the number of target chromophores increases and the depth increases, the inverse problem becomes ill-posed, underdetermined and noise-sensitive, further compounding inaccuracies. Finally, overlapping spectra with similar absorption patterns introduce inaccuracies to the linear model calculations. Overall, conventional linear unmixing methods, such as non-negative least squares (NNLS), frequently misinterpret the endogenous spectra, leading to quantification errors or the “hallucination” of molecules.

To overcome these limitations, we introduce **photoacoustic fingerprinting (PAF),** a machine-learning framework that redefines the spectral unmixing problem as a nonlinear mapping between spectral shape and molecular composition, treating the inversion effectively as a pattern recognition task. Unlike inversion-based methods that attempt to explicitly correct for fluence, PAF implicitly learns the joint effects of fluence, noise, and spectral overlap through training, enabling robust inference without requiring explicit modeling. This approach is conceptually analogous to magnetic resonance fingerprinting, which transformed MRI by replacing parameter-specific inversion with pattern recognition-based quantification [13]. However, unlike MRI, where acquisition can be explicitly designed to encode parameters, photoacoustic signals are shaped by unknown and spatially varying optical fluence, making direct inversion fundamentally unstable. This perspective fundamentally shifts photoacoustic unmixing from a strictly physics-constrained inversion toward a data-driven chemical fingerprinting paradigm while still benefiting from physics informed priors, analogous to spectroscopic identification in analytical chemistry. We define the *photoacoustic fingerprint* as the vector of signal amplitudes across an oversampled wavelength set, where we treat the sequence of multispectral signal amplitudes not as a linear sum of pure components, but as a complex “fingerprint” encoding both molecular concentration and spectral distortion. Our work is inspired by recent trends in the integration of machine learning with PAI and inverse problems [14–16]. Recent work from our group has demonstrated the effectiveness of deep learning for solving inverse problems in photoacoustic imaging, including reconstruction from undersampled data and physics informed priors [17, 18]. In spectroscopy, data driven models can implicitly learn complex wavelength dependent variations and noise statistics, enabling more robust spectral unmixing under realistic imaging conditions [19]. Complementary advances in detection sensitivity and speed, such as dual-channel photoacoustic microscopy, further improve fidelity and SNR, reinforcing the need for robust computational frameworks capable of extracting accurate molecular information from increasingly high-quality yet complex datasets [20].

This method resulted in a higher reconstruction accuracy of molecular concentrations in synthetic data across various signal-to-noise ratios (SNR), as well as the ability to identify components in phantoms with multiple chromophores, some with overlapping absorption spectra. Then, we demonstrated PAF’s capability in analyzing experimentally acquired data. To validate the framework in biological tissue, we first apply PAF to *ex vivo* mouse liver samples from control and high-fat diet models, demonstrating the ability to detect steatosis-induced lipid accumulation. Lastly, we extend this to a clinical setting by imaging human arteries extracted from patients with peripheral artery disease. In vascular disease, where plaque composition and thrombus organization critically influence outcomes, the inability to resolve overlapping molecular signatures limits the diagnostic utility of current imaging methods. PAF successfully resolved plaque composition and thrombus organization. Together, these results position PAF as a framework for translating photoacoustic imaging from qualitative contrast mapping toward quantitative, clinically actionable molecular diagnostics.

## 2. Theory of Photoacoustic Fingerprinting

The photoacoustic signal arises from the thermo-elastic expansion of tissue following pulsed optical absorption [3, 21], and can be modeled as a function of the local optical fluence, concentration and absorption spectrum of a sample. The linear unmixing method assumes the following model:

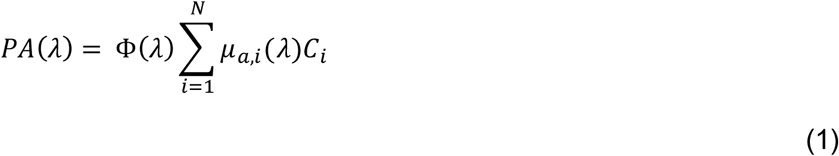

Here *PA*(*λ_i_*) is the reconstructed PA signal at a specific wavelength λ, Φ(λ) is the local, wavelength-dependent optical fluence, *β,_a,i_* is the absorption coefficient spectrum per chromophore *i* and *C_i_* is its concentration. We define the **PA fingerprint** as the vector of PA signals at an oversampled wavelength grid.

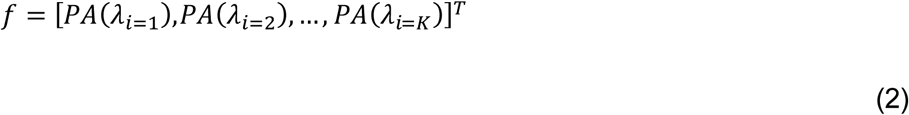

Here, K (number of sampled wavelengths) is significantly greater than N (number of chromophores), creating an overdetermined system that allows for enhanced noise resilience and spectral feature extraction. This high-dimensional representation captures subtle spectral features that help disambiguate molecules with overlapping absorption peaks.

The conventional linear unmixing models assumes Φ(λ) either is known and uniform (unit fluence) or can be removed exactly by energy normalization. In practice Φ(λ) varies with depth and tissue optical properties (absorption and scattering), and therefore acts as a multiplicative spectral distortion that can bias linear inversions. To mitigate fluence variation effects, the raw PA signals are commonly normalized by the measured or modeled incident light energy [2, 22]. The linear model can be solved using linear matrix inversion, with or without non-negative constraints. Equation 1 represents the forward model for the RNN, where the goal is to invert and calculate *C_n_* given the spectral fingerprint. RNNs are particularly well suited for sequential or ordered data, allowing the model to exploit wavelength-to-wavelength correlations that improve robustness and accuracy over conventional linear unmixing [16]. The objective of PAF is thus to invert ***f*** for the concentration vector [*C*_1_, …, *C_N_*], producing a unique mapping between measured PA fingerprints and their underlying molecular compositions.

The method used for comparison is *non-negative least squares* (NNLS) matrix calculations, as real-world concentrations will be non-negative. Figure 1b demonstrates how increasing the number of sampled wavelengths in a synthetic data set decreases observed NNLS error. Because the fingerprint encodes complex noise-perturbed absorption features across a large number of wavelengths, we employ a recurrent neural network to learn the nonlinear mapping from spectral fingerprints to concentration vectors, thereby improving robustness over conventional linear unmixing.

**Figure 1:**
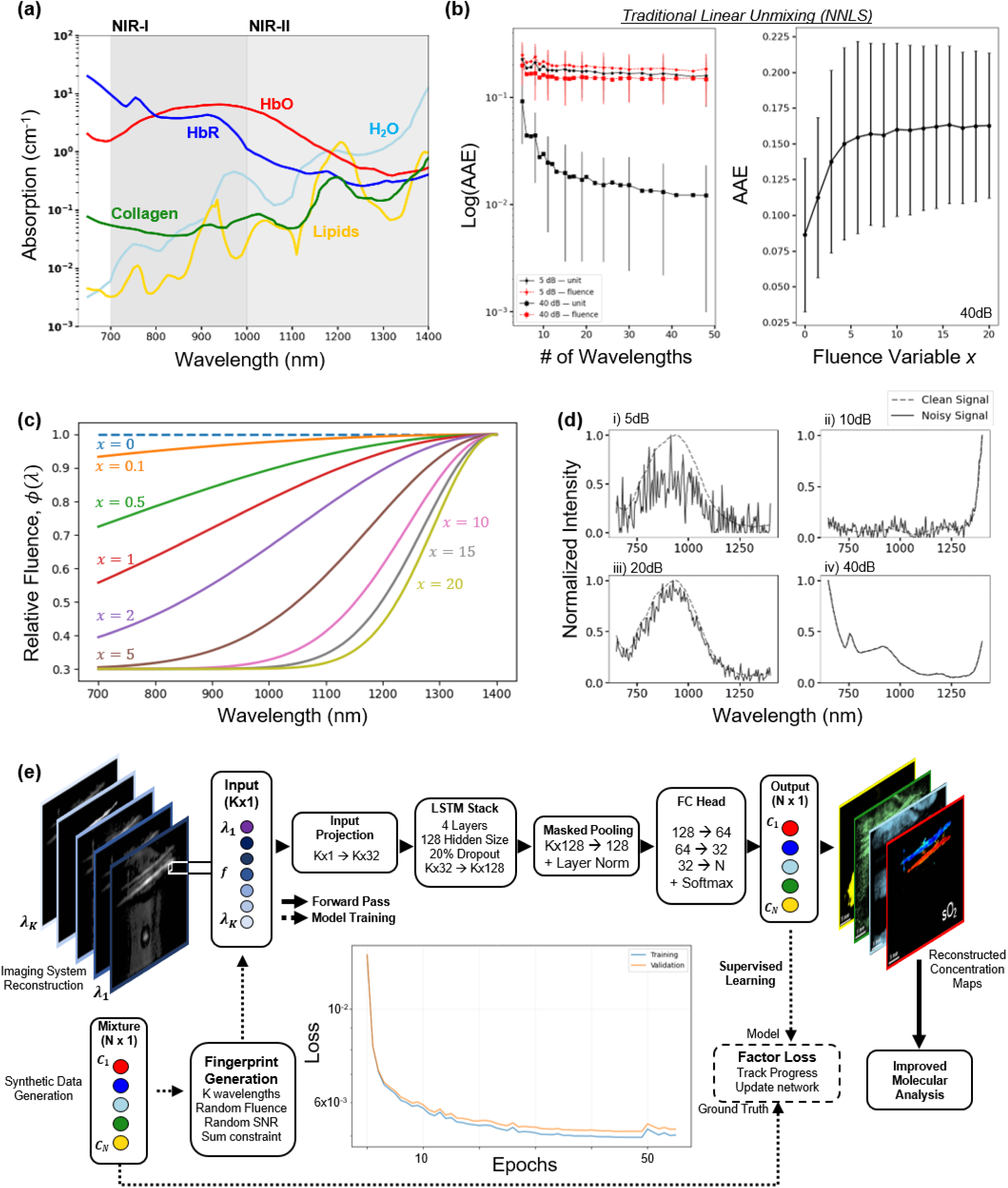
Working principles of PAF. **(a)** Pure absorption coefficient spectra used to build the spectral library: HbO₂, HbR, lipids, water and collagen. **(b)** NNLS average absolute error (AAE) as a function of the number of sampled wavelengths under low (5 dB) and high (40 dB) SNR, with and without fluence distortion. For a 40dB signal, increases the severity of the fluence variable increases error. AAE decreases with more wavelengths and higher SNR with unit fluence, illustrating the value of spectral oversampling. **(c)** Example fluence profiles produced by the parametric Beer’s law prior (fluence parameter varied). **(d)** Example synthetic photoacoustic fingerprints at varying SNR levels (5–40 dB), demonstrating spectral variability used for model training. Under unit fluence (solid black lines), increasing the number of wavelengths reduces error, consistent with overdetermined inversion. In contrast, under wavelength-dependent fluence distortion (dashed red lines), error plateaus despite increasing spectral sampling. **(e)** Overview of the PAF framework. Multispectral photoacoustic signals are treated as high-dimensional fingerprints and processed by a RNN to infer molecular concentrations. The architecture consists of an encoder, stacked LSTM layers, masked pooling, and a fully connected readout. Training is performed using synthetic fingerprints spanning a wide range of noise, fluence, and mixture conditions.

## 3. Materials and Methods

### 3.1 Endogenous Agents – Photoacoustic Spectral Library

Many endogenous chromophores in tissue naturally provide contrast for photoacoustic imaging. Each chromophore has a unique absorption profile across wavelengths, useful for identifying structures and functions. These agents are beneficial for their biocompatibility, endogenous abundance, and minimal safety concerns [23]. Key endogenous contrasts in PAI include hemoglobin (oxygenated HbO_2_ and deoxygenated HbR) [24, 25], DNA/RNA [26], melanin [27], lipids [28, 29], water [30], glucose [31], and collagen [32].

We defined the **photoacoustic spectral library** as a chemically motivated reference set derived from pure absorption coefficient spectra. This approach mirrors classical spectroscopy library construction, allowing PAF to operate as a generalized chemical classifier and quantifier rather than a fixed unmixing solver. Figure 1a shows the assembled spectral library of five dominating endogenous chromophores in the near infrared (NIR) wavelength range [33–36]. All spectra are linearly interpolated onto a common wavelength grid based on the signal (700-970nm) and idle (1160-1400nm) ranges of the lasers, retaining their raw absorbance units for network training.

### 3.2 Synthetic Data Generation

Robust training of the PAF model requires a diverse dataset that captures the variability encountered in biological tissues. To achieve this, we generated a synthetic dataset of 1 million spectral sequences. Each sequence was constructed by sampling concentrations of hemoglobin, lipids, collagen and water. Each concentration vector was normalized such that the sum of all components equals one. Although simplified, this synthetic framework enables controlled exploration of failure modes that cannot be isolated in experimental measurements, where ground truth is generally unknown.

Half the vectors were sampled equally across five mixture types designed to represent varying spectral complexity:

- **Pure**: one chromophore at 100%
- **Extreme Mixtures**: one chromophore at 90-99%, others sharing the remainder randomly
- **Dominant Mixtures**: one chromophore at 70-90%, others sharing the remainder randomly
- **Random Ratios**: all concentrations drawn randomly
- **Binary Mixtures**: exactly 2 chromophores, each >1%

In addition to spectral mixtures, half the fingerprints were generated across representative anatomical patters to mimic realistic biological composition:

- **Vessel-like**: hemoglobin dominant with oxygenation variations
- **Fat-dominant** and **collagen-rich** regions
- **Background tissue:** primarily water-based absorption

This design ensured that the model was exposed to a wide range of spectral and physiological scenarios, improving generalization across tissue types and biological contexts.

The signal at K sampled wavelengths is modeled as

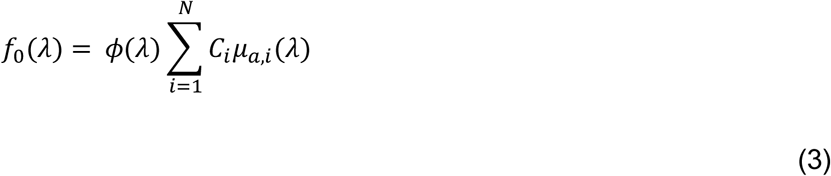

For photoacoustic imaging, fluence decay is governed by the diffusion approximation and follows more complex wavelength-dependent absorption and scattering. To generate wavelength-dependent distortions during training we applied a simple exponential attenuation model (Beer’s Law) as a parametric fluence prior:

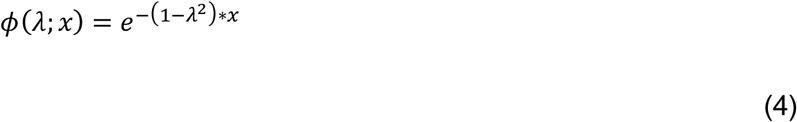

Where *x* is a scalar fluence severity parameter sampled from [0, 20]. The range x∈[0, 20] was empirically chosen to span a broad dynamic range of attenuation strengths, from near-uniform fluence to severe spectral distortion, ensuring sufficient diversity in training while remaining within the numerically stable bounds of the exponential model. This 1D exponential model captures basic wavelength dependence (shorter wavelengths being attenuated more strongly than the longer wavelengths) and enables controlled experiments probing PAF’s robustness to multiplicative spectral distortion. Importantly, the 1D Beer’s law ignores the lateral heterogeneity. Therefore, the synthetic fluence used in training represents a convenient prior that partially approximates typical depth-dependent spectral reddening but does not replicate accurate light transport in turbid media. Rather than explicitly correcting for fluence, this approach allows the model to learn a statistical mapping between distorted spectral fingerprints and underlying molecular composition.

To simulate realistic noise conditions, we independently drew a signal-to-noise ratio (SNR) in decibels from a uniform distribution over [3, 40] dB for each fingerprint. Gaussian distributed noise with zero mean and unity power is added to each fingerprint. We define the signal power and noise power as:

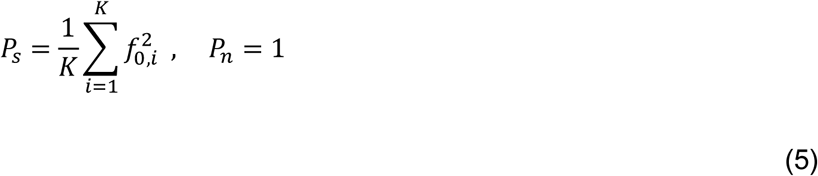

and the desired SNR in decibels as:

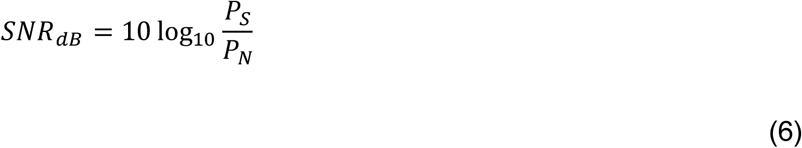

We can solve the scaling factor α,

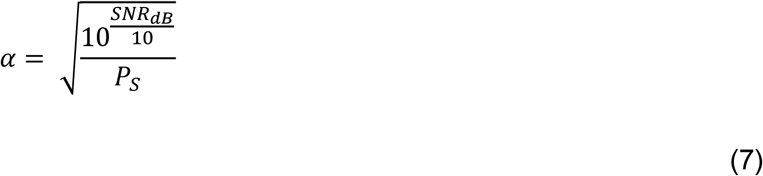

The final noisy fingerprint is then constructed as

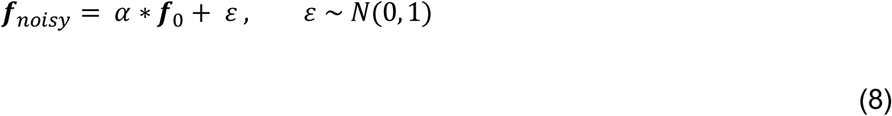

Wavelength-dependent relative fluence variations and wavelength-independent Gaussian noise addition mimics the fingerprints behavior observed in real measurements, allowing the synthetic data generation to be physics driven. Figure 1d illustrates randomly sampled fingerprints at 5, 10, 20 and 40dB SNR. Each fingerprint was normalized by its maximum value to emphasize spectral shape in training. By spanning a broad SNR range and including multiple mixture and anatomical classes, the training dataset captured both physical and biological variability, enabling PAF to perform robust spectral unmixing across realistic conditions.

These simulated fingerprints serve as the forward-model training for the RNNs described in Section 4.3. By exposing the network to diverse spectral patterns and noise regimes, the model learns to generalize across spectral variations and complex mixtures, rather than memorizing idealized spectra.

### 3.3 Recurrent Neural Network (RNN) Architecture, Training, and Performance Markers

The RNN serves as a function approximator for the fingerprint-to-concentration mapping, rather than being the primary source of novelty in this work. All modeling and reconstruction work for PAF and NNLS was performed in Python 3.12.5 using PyTorch 2.7.1 on a desktop equipped with an NVIDIA GeForce RTX 4060 Ti GPU (8 GB VRAM) and 32 GB system RAM. The entire dataset consists of synthetic fingerprints. From 1 million total samples, 80% were used for training and 20% held out for validation during training. Unique concentration test sets of size 50k were generated for further model validation. Each noisy fingerprint vector was normalized by dividing by its maximum value, ensuring inputs span [0, 1], putting emphasis on spectral shape for quantitative discovery rather than overall intensity. Because the spectral data represents ordered wavelength-dependent sequences, we implemented a RNN to exploit spectral correlations and contextual dependencies between adjacent wavelengths. The final architecture (Figure 1e) consisted of the input fingerprint (a vector of length K,*f* = [*PA*(λ_*i*=1_),*PA*(λ_*i*=2_), …, *PA*(λ_*i*=*K*_)]^*T*^), an encoder, and readout head (a linear layer mapping the final hidden state to an output vector of size N (one per chromophore trained), followed by a softmax activator to enforce 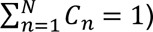.

The following hyperparameters, selected through grid search to balance performance and computational cost, were used for training: Batch training size = 32, learning rate = 1e-4, epochs = 100. Training utilized the AdamW optimizer (learning rate, α = 1e-4, β = [0.9, 0.999], weight decay = 1e-5) algorithm to calculate gradients and update weights. Smooth L1 Loss (Huber loss) was used to calculate the loss between predicted concentrations and the ground truth. After training, the model state was saved and subsequently loaded for test set evaluation. It is important to note that pure ratios of molecules were still generated in testing sets at various fluence and SNR values.

Three models were trained for synthetic validation. One model was exactly-determined with unit fluence, where the number of wavelengths equals the number of chromophores. One model was over-determined with unit fluence, with the number of wavelengths much greater than the number of chromophores. Finally, to compare the effects of optical fluence, a third model was trained on an over-determined set of wavelengths and random fluence in fingerprint generation.

Although training was optimized using factor loss, the overall model performance across all SNRs was evaluated using the following quantitative metrics, as shown in the Results and Discussion section: Coefficient of determination (R^2^), Root-mean-square-error (RMSE), and Slope of predicted concentrations vs. true concentrations. These metrics were analyzed through averaging across different test sets. Furthermore, average absolute error (AAE) for each model’s predicted concentration vectors were calculated, ensuring a robust way to track molecular reconstruction against SNR.

### 3.4 Phantom and Ex vivo Liver and Human Artery Sample Preparation

After validating the model on synthetic data, we increased experimental complexity using a controlled phantom study, allowing us to test PAF against physical noise and system-specific artifacts. Then, to evaluate the model in a realistic biological environment where scattering and absorption are non-uniform, we conducted *ex vivo* imaging on mouse livers and human arterial tissues. The phantoms, biological tissues, and human arteries were imaged using the Diffractive Acoustic Tomography (DAT) system developed in our lab [37]. The reconstructed datasets span a wide range of distinct photoacoustic signal regimes. The multi-component phantom provides a known chromophore mixture reconstruction, while the *ex vivo* biological tissues and clinically relevant human samples provide a realistic tissue analysis. This allows for a comprehensive assessment of PAF’s reconstruction robustness and molecular specificity across increasing experimental complexity.

For experimental datasets, we corrected for pulse-to-pulse laser energy using a calibrated power meter (Ophir). This energy normalization removes per-pulse laser energy fluctuations but does not compensate for depth- or tissue-dependent fluence variations. For PAF training and inference we also applied per-voxel max normalization (each fingerprint scaled to [0,1]) to emphasize spectral *shape* rather than absolute amplitude. NNLS comparisons use raw energy-normalized fingerprints (no max-scaling), because NNLS relies on relative amplitudes to estimate concentrations. We verified that NNLS results were largely insensitive to secondary max-scaling, but the normalization choices directly affect each method’s sensitivity to fluence-driven dynamic range changes and should be considered when interpreting quantitative outputs. NNLS received the same spectral library used for PAF model training for concentration vector reconstruction. From Python’s Scipy Optimized package, the NNLS function was used to calculate the concentration vector given the spectral library and measured PA signals. Both models output concentration vectors including HbO, HbR, lipids, collagen and water. Finally, the absolute hemoglobin concentrations are used to calculate oxygen saturation:

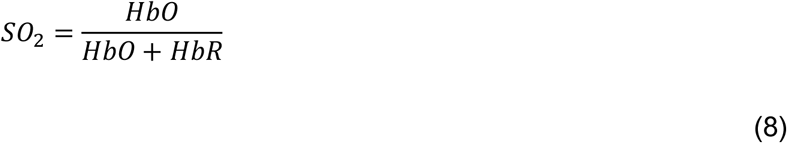

#### 3.4.1 Multicomponent Phantom Preparation

The multicomponent phantom was designed to represent known soft tissue compositions containing multiple absorbers in the detected PA signals. It consisted of five key components: Oxygenated bovine blood (prepared with sodium bicarbonate), deoxygenated bovine blood (prepared with sodium hydrosulfide), melted butter to model lipid-rich regions, a droplet of collagen mixed with water, and a drop of water.

All samples were placed on a transparent membrane with water beneath to provide acoustic coupling and imaged using the DAT system [37]. A Surelite Continuum II-10 pump laser and Surelite OPO Plus were used for PA illumination, providing volumetric photoacoustic imaging over the 700-970 nm and 1160-1400 nm wavelength ranges. Heavy water was used for the acoustic wave coupling, minimizing near-infrared absorption. The reconstructed PA images were analyzed using NNLS and PAF to directly compare unmixing performance across all five chromophores.

#### 3.4.2 Ex vivo Fatty Liver Preparation

To further assess PAF’s performance in biologically relevant tissues, we applied the model to *ex vivo* mouse livers. Two male albino B6 mice were used for this study, under protocols approved by the Duke University IACUC (animal protocol 60681). One mouse was maintained on a control diet and the other on a high fat diet (HFD) for six months, a regimen known to induce hepatic steatosis (fatty liver disease) [38]. The livers were excised and imaged by DAT. Imaging was conducted from 700-970nm and 1160-1400nm at 10 nm steps. This experiment served to evaluate PAF for lipid accumulation in metabolically altered tissue.

#### 3.4.3 Ex vivo Human Artery Sample Preparation

To demonstrate PAF’s ability to analyze clinical samples, *ex vivo* human artery section samples were obtained post-operatively from above-knee and below-knee amputations performed on patients with known peripheral artery disease (PAD), under protocols approve by the Duke University Institutional Review Board (Duke IRB#: Pro00065709). The arteries were stored in phosphate buffered saline (PBS) at 4°C before imaging. The reconstructed datasets were analyzed using PAF to identify molecular distributions associated with vascular plaque formation. A separate artery was analyzed using scanning electron microscopy to provide an individual comparison.

## 5 Results and Discussion

### 5.1 Synthetic Reconstruction Comparison

We first sought to test whether PAF can overcome the fundamental limitations of inversion-based unmixing under controlled but realistic conditions where NNLS fails. Unknown fluence and variable SNR represent the dominant sources of error in practical photoacoustic imaging. Figures 2a-b summarize the parity plots of true vs. predicted concentrations under two sampling regimes: the exactly-determined case (Figure 2a), and the overdetermined case (Figure 2b). In both cases, data points are color-coded by SNR to illustrate reconstruction behavior across noise levels. Figure 2c summarizes the parity plots for the over-determined case with fluence variations, colored by the fluence variable. Importantly, these conditions directly reflect the dominant sources of error in clinical photoacoustic imaging, where fluence and SNR are unknown. While synthetic data cannot fully capture tissue complexity, it provides controlled access to ground truth, enabling systematic evaluation of failure modes that are otherwise unobservable in experimental data. Test sets were generated independently from training distributions to evaluate generalization across unseen mixtures and noise realizations. For statistical analysis, the coefficient of determination (R^2^), root-mean-square error (RMSE), and slope were calculated for the models of five unique testing sets. Average absolute error was calculated across SNR per chromophore. A two-tailed t-test was used to calculate p-values between the PAF and NNLS overdetermined cohorts.

**Figure 2:**
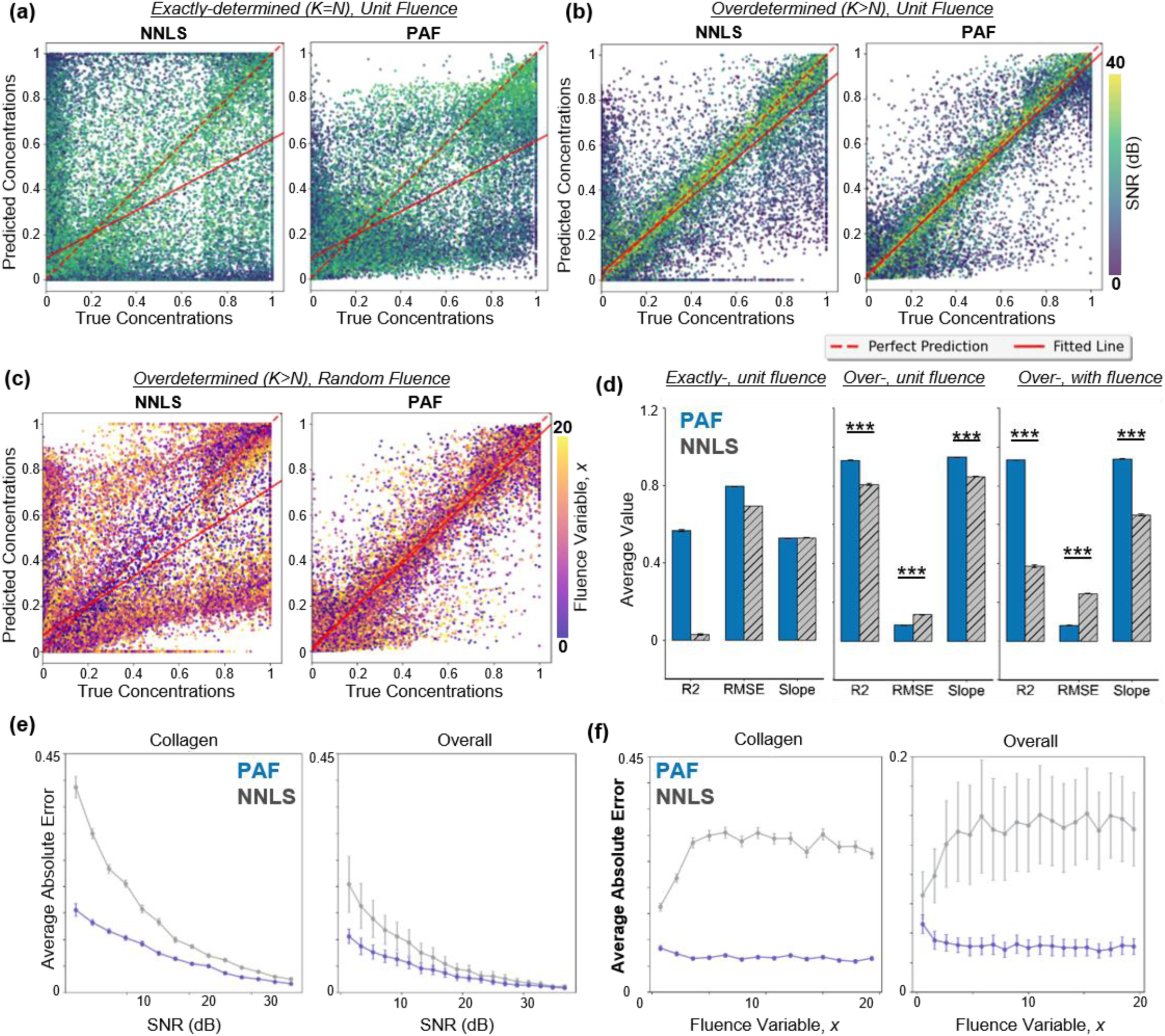
Synthetic reconstruction comparisons between NNLS and PAF. (a) Exactly-determined case (K=N): predicted vs true concentrations for NNLS (left) and PAF (right). Both methods exhibit substantial variability due to sensitivity to noise and spectral overlap. (b) Over-determined case (K≫N): both methods improve, but PAF shows tighter agreement with ground truth. (c) Over-determined case with randomized fluence: PAF maintains accuracy while NNLS degrades substantially. (d) Summary metrics (R², RMSE, slope) showing PAF’s consistent improvements with uniform fluence and random fluence. (e) AAE vs SNR for collagen (spectrally challenging) and all chromophores in the over-determined, unit fluence case: PAF yields lower error and variance, especially at low SNRs. (f) AAE vs fluence in over-determined, uncertain fluence case: NNLS error rises with fluence uncertainty while PAF remains relatively flat.

In the exactly-determined casewith unit fluence (Figure 2a), NNLS and PAF showedlarge scatter and systematic under- and over-estimation across all concentration levels. In the over-determined case with unit fluence (Figure 2b), where the number of wavelengths far exceeds the number of chromophores, both methods improved due to increased spectral sampling. Furthermore, PAF outperforms NNLS with a higher *R*^2^ (0.931±0.0008 vs. 0.807±0.0047), improved RMSE (0.081±0.0004 vs. 0.136±0.0017), and a slope closer to unity (0.947±0.0049 vs. 0.848±0.0027), significantly increasing reconstruction accuracy (Figure 2d). These results confirm that PAF remains generalizable in low-SNR regimes, increasing quantitative accuracy higher than NNLS when the system becomes overdetermined. When fluence variation was added to the fingerprints, NNLS dropped in accuracy significantly, while PAF was able to learn the non-linear relationships to unmix the concentrations (Figures 2c and 2d). With fluence introduced, PAF outperformed NNLS with a higher *R*^2^ (0.933±0.0016 vs. 0.385±0.0083), improved RMSE (0.080±0.0011 vs. 0.244±0.0014), and a slope closer to unity (0.934±0.0017 vs. 0.650±0.00248). PAF substantially outperformed NNLS in scenarios where wavelength-dependent fluence variations are present, demonstrating that spectral sampling and distortions affect accurate spectral unmixing.

Figure 2e further quantified the over-determined unit fluence model’s performance through the average absolute error (AAE) as a function of SNR. Collagen, the most spectrally challenging chromophore due to its overlap with lipid and water absorption, exhibited the largest improvement. PAF reduced error up to 50% at low-SNRs as compared to NNLS. The overall average error across all chromophores follows a similar trend: PAF consistently yields lower error and reduced variance, particularly below 20 dB, demonstrating stronger noise resilience. When looking at the fluence variation model, the largest improvements are again for collagen. Plotting the AAE as a function of the fluence variable, *x*, shows that as fluence increases, NNLS performance goes down. In contrast, the PAF performance across all fluence variables is nearly consistent (Figure 2f). Together, the synthetic test sets highlight PAF’s robustness to both noise and fluence.

Beyond specific chromophore reconstruction, Table 1 summarizes the accuracy across mixtures in the overdetermined regime. PAF improved *R*^2^ across all mixtures, most notably in dominant and random mixtures, where concentration variability was highest.

**Table 1:** R^2^ Values for Mixture Groups by Model.

|  | Pure | Extreme | Binary | Dominant | Random |
| --- | --- | --- | --- | --- | --- |
| **NNLS** | 0.888±0.0044 | 0.867±0.0061 | 0.774±0.0056 | 0.791±0.0053 | 0.295±0.0185 |
| **PAF** | 0.981±0.0014 | 0.981±0.0012 | 0.869±0.0044 | 0.946±0.0023 | 0.644±0.0044 |
| **Total Number** | 7500 | 10015 | 12505 | 10670 | 9310 |

These results indicate that the RNN-based PAF model more effectively captured the non-linear relationships between wavelength-dependent features and concentration ratios than NNLS, which assumes linear relationships among spectral components. The improvement is most evident in complex-mixture and collagen-rich regimes.

Our synthetic results highlighted three key advantages of the PAF approach: Robustness to noise and fluence, flexibility for complex mixtures, and high sensitivity to collagen samples with low SNRs. By training the model focusing on spectral shape rather than signal intensity, PAF maintains reliable quantitative accuracy. This is particularly beneficial for low-SNR multi-component mixtures or deep tissue imaging where simple linear inversion often fails to disentangle overlapping spectral features or correct for the fluence variances. In other words, when applied to experimental PA measurements, where fluence is only partially accounted for with recorded laser energy and system noise is unknown, PAF’s data-driven nature promises more reliable unmixing than physics-only models. The improved slope indicates less systematic under- and over-estimation across the full concentration range, especially in low SNR imaging. Maintaining a consistent response across SNR levels is essential for reliable molecular quantification for deep-tissue molecular imaging. Together, these results establish that robustness to noise and fluence, on top of increased spectral sampling, is the key requirement for accurate molecular quantification

### 5.2 Multicomponent Phantom Reconstruction

Next, we applied PAF to phantom data to evaluate performance in a realistic imaging context. We constructed a phantom comprising bovine blood, butter, bovine collagen, and water (Figure 3a). The phantom was imaged on our custom-built DAT system with a measured SNR around 10dB. Three-dimension PA volumes were reconstructed and compressed via maximum amplitude projection (MAP) at each wavelength to yield 2D multispectral stacks.

**Figure 3:**
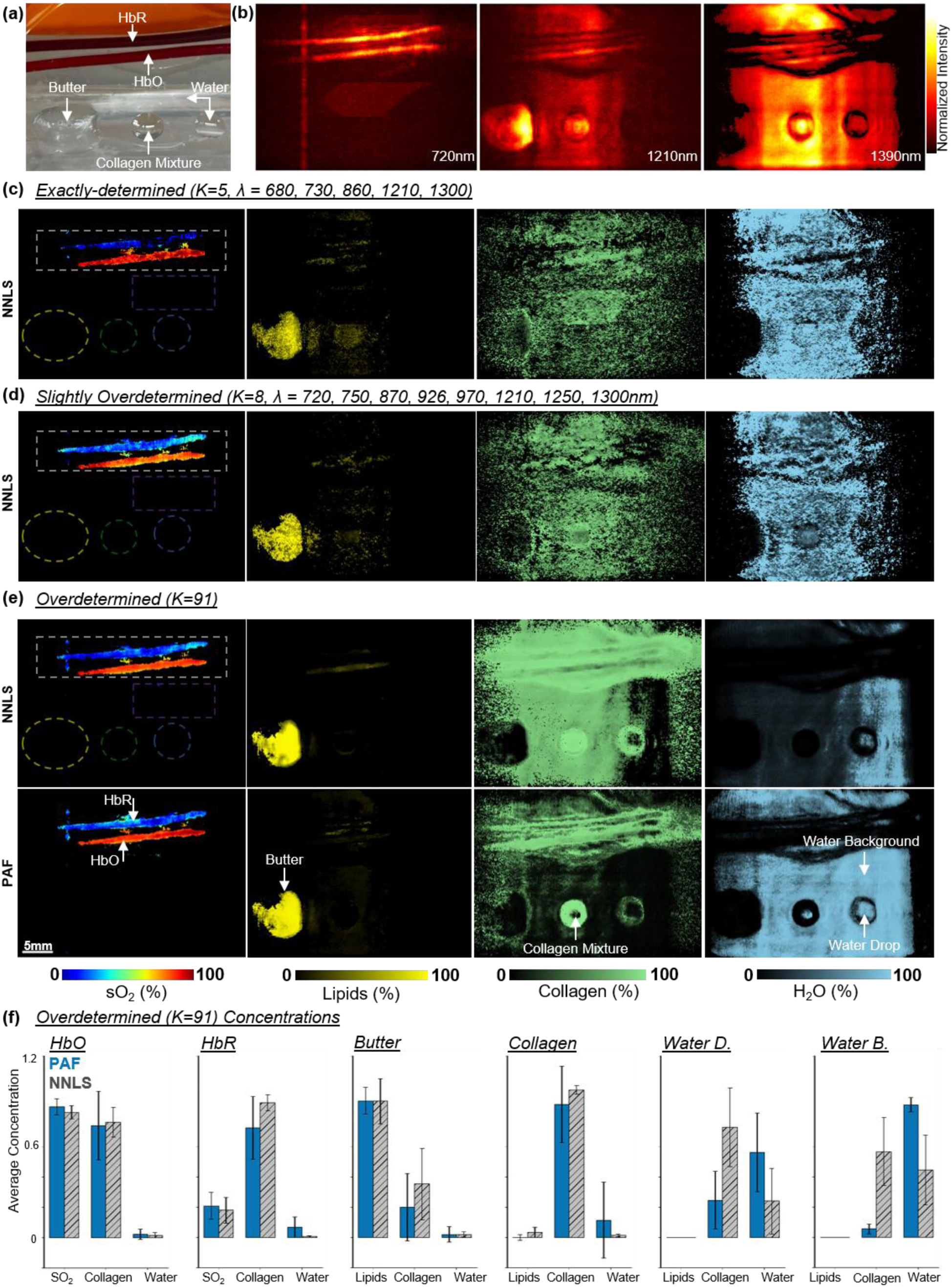
Multicomponent phantom imaging. **(a)** Top-down photo of the phantom and ROIs, including blood tubes, melted butter, collagen, water droplet, and background coupling. **(b)** Representative MAPs at 720nm, 1210nm, and 1390nm, illustrating wavelength-dependent contrast across chromophores. Reconstructed maps for sO₂, lipids, collagen and water using **(c)** an exactly determined set wavelengths, **(d)** a slightly-overdetermined set of wavelengths, and **(e)** an overdetermined set of wavelengths. In the overdetermined case with low-SNR conditions (∼10 dB), NNLS overestimates collagen in background regions while PAF shows improved chemical specificity as it localized collagen and recovered water contrast more accurately. **(f)** Quantified concentrations within ROIs (mean ± s.d.), across multiple chromophores.

Colored areas in and Figure 3c highlight the bovine blood tubes (white), butter (yellow), collagen (green), the water droplet (blue), and some water background (purple). We applied both PAF and NNLS unmixing to the multispectral PA images depending on the number of sampled wavelengths and quantified the concentration maps for the components of the overdetermined group (Figure 3f).

For NNLS, as the number of wavelengths used for reconstruction increased, it is clear to see the linear model begin to localize the chromophores more accurately. The overdetermined set performed the best, where NNLS and PAF reconstructed similar sO_2_ and lipid concentrations. However, substantial differences emerged for collagen and water, the two chromophores with the most overlapping absorption spectra in the NIR wavelength range. NNLS overestimated collagen in the system and in the background. In contrast, PAF localized collagen more sharply and with lower background abnormalities. In addition, PAF better captured the water droplet and the coupling background, while NNLS underestimated water in both regions. This demonstrated that PAF’s data-driven spectral mapping successfully disentangles overlapping spectra even under the challenging low-SNR conditions (∼10 dB). Nevertheless, both PAF and NNLS notably exhibited collagen overestimation within the blood tubes, likely due to the spectral crosstalk between collagen and water or the unknown absorption addition of the silicon tubes to the photoacoustic signal.

Interestingly, both models achieved similar spectral reconstructions (Figure 4) matching the overall shape of the multispectral fingerprint. As shown previously in Figure 3, NNLS can attribute water rich regions as collagen. In contrast, PAF can better confine collagen to its true location. This reveals an important limitation of linear unmixing metrics, where outcome spectral fidelity does not necessarily reflect chemical accuracy. This result highlights a fundamental limitation of conventional unmixing: *accurate spectral reconstruction does not imply accurate molecular interpretation*. In contrast, PAF is not merely fitting spectra, but performing meaningful chemical classification, a distinction that is critical for *in vivo* molecular pathology and tissue composition analysis. By learning the nonlinear relationship between spectral features and molecular composition during training, PAF avoided the systematic unmixing errors that NNLS exhibits despite achieving similar spectral reconstruction accuracy.

**Figure 4:**
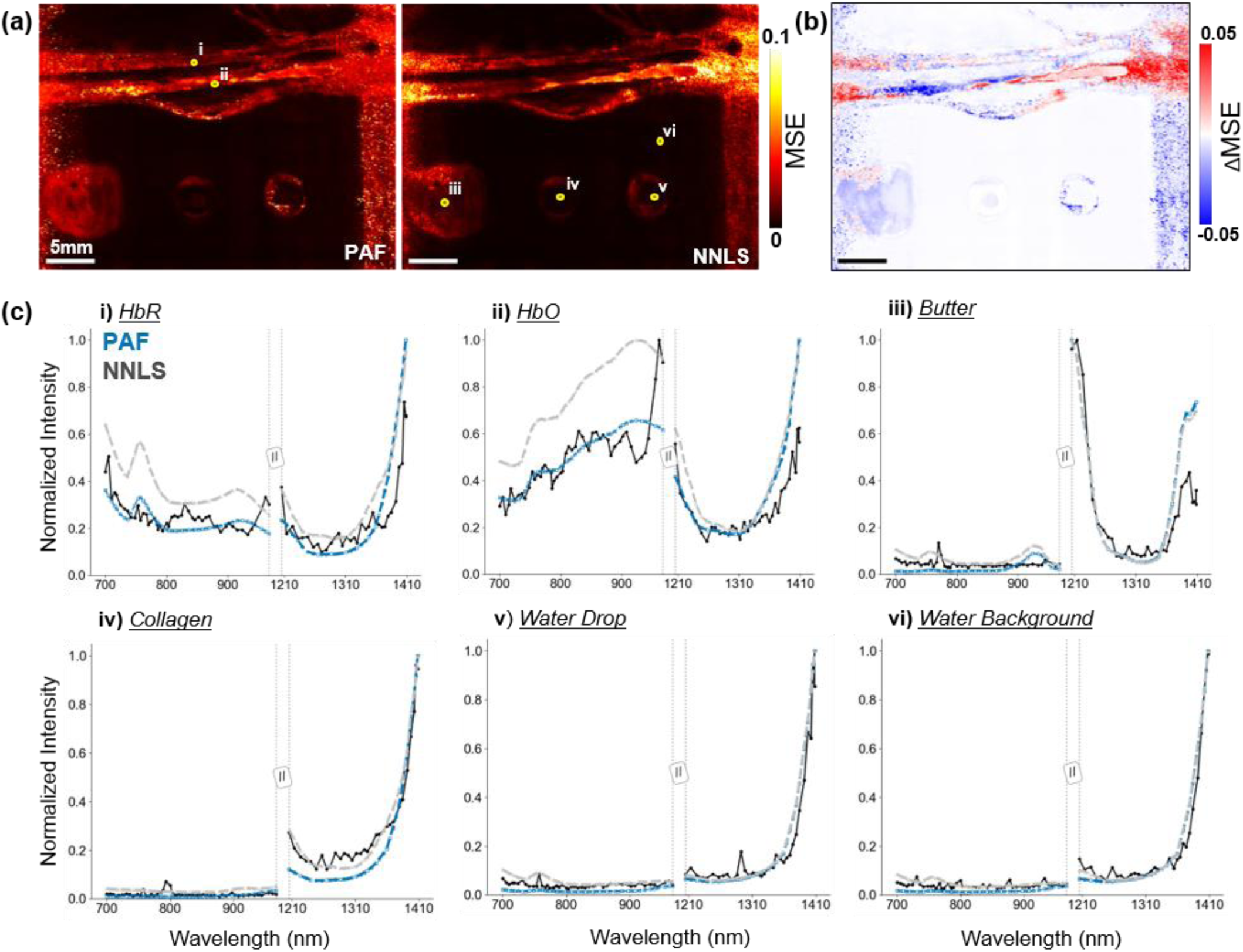
Spectral fidelity and comparison between PAF and NNLS. **(a)** Mean squared error (MSE) maps. Darker regions indicate lower MSE. **(b)** Difference map (PAF − NNLS) highlighting locations with larger reconstruction discrepancies. **(c, i–vi)** Example measured fingerprints and model reconstructions from regions corresponding to HbR, HbO, background, water drop, collagen, and butter (lipid).

### 5.3 Ex vivo Fatty Liver Analysis

To extend PAF validation beyond controlled phantoms and assess its sensitivity to biological variation, we applied the model to *ex vivo* mouse livers obtained from control and HFD cohorts. The liver serves as an ideal model for evaluating lipid content as chronic exposure to HFD induces lipid accumulation and steatosis, producing reliable chemical differences in tissue composition. Figure 5 shows the unmixed lipid and collagen concentration maps for both liver samples along with corresponding lipid signal histograms. The control liver exhibited low lipid signals, consistent with expected hepatic composition. In contrast, the HFD liver displayed elevated lipid accumulation, indicative of fatty liver disease. In this study, histogram analysis revealed clear shift in the HFD liver, with mean lipid concentration increased by 2.6-fold, when compared to the control liver.

**Figure 5:**
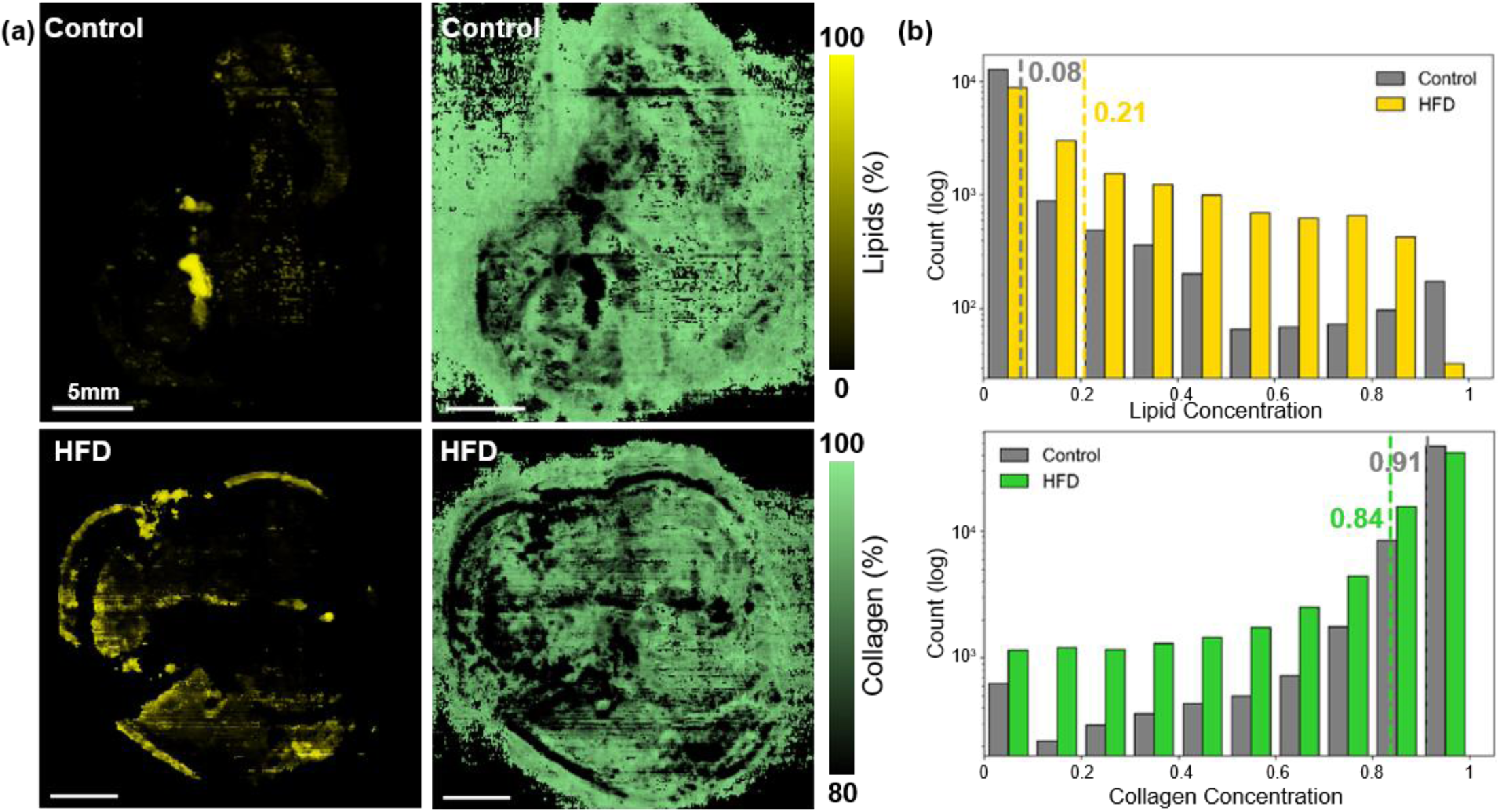
Ex vivo mouse liver results (control vs high-fat diet). **(a)** PAF reconstructed lipid and collagen maps for control (top) and HFD (bottom) livers. **(b)** Lipid and collagen distributions quantified from histograms. The HFD sample shows a rightward shift in lipid content and a relative reduction in collagen signal, indicating increased lipid accumulation and compositional change. Mean lipid levels increase from 0.08 (control) to 0.21 (HFD), while collagen distributions show an opposing trend.

Although limited to a pilot comparison, these proof-of-concept results aligned with the expected biochemical changes associated with fatty-liver development and demonstrate that PAF can detect and quantify physiologically relevant lipid variations without the need for exogenous labelling. PAF preserved biologically meaningful contrast consistent with known disease pathology. Importantly, this is achieved without explicit modeling of fluence or tissue optical properties. In the *ex vivo* setting, extended wavelength ranges, high spectral sampling, and aggressive signal averaging are feasible, all of which can further improve PAF’s accuracy.

### 5.4 Ex vivo Human Artery Sample Analysis

To evaluate clinical relevance, we applied PAF to *ex vivo* human arteries from patients with peripheral artery disease (PAD), where resolving plaque composition and thrombus organization remains a major unmet need. Artery sections were excised *ex vivo* from patients undergoing above- and below-knee amputations due to peripheral artery disease (PAD), a condition characterized by plaque buildup and thrombus formation that obstructs blood flow [39–41]. PAD is not only a disease of luminal narrowing due to atherosclerotic plaque, but also frequently involves extensive arterial calcification [42–44].

Although non-contrast CT is highly sensitive to vascular calcium and can quantify overall calcific burden, it provides no information about non-calcified plaque components such as lipids, organized thrombus, fibrous remodeling, extracellular matrix content, or oxygenation state. These features are central to PAD progression, procedural success, and prognosis, yet remain invisible on CT. Therefore, combining a CT-identified calcification with PAF analysis of the excised artery provides a powerful bridge between anatomical imaging and molecular characterization.

Clinical sagittal-view CT images of a patient’s right leg were reviewed to confirm calcification (Figure 6a). Following the above-knee amputation, a segment of the popliteal artery containing suspected plaque and thrombus was excised (sample VGF271). The reconstructed multispectral PA images were analyzed using PAF to generate quantitative maps of sO₂, lipids, collagen, and water (not shown). Figures 6d-h present the PAF maps for the artery segment, showing clear biochemical distributions. Due to the destructive nature of fixation required for scanning electron microscopy (SEM) and histology, co-registration with the same specimen is not feasible without altering lipid content. Therefore, validation was performed on closely matched samples exhibiting similar pathological features. A separate segmented human artery (sample VGF279) was imaged using SEM to reveal plaque buildup and a packed RBC region (Figure 6j-l). Despite the absence of direct co-registration, the spatial patterns observed with PAF are consistent with known morphological and biochemical features of thrombus and plaque. These findings suggest that PAF can provide biochemical information inaccessible to conventional clinical imaging modalities such as CT

**Figure 6:**
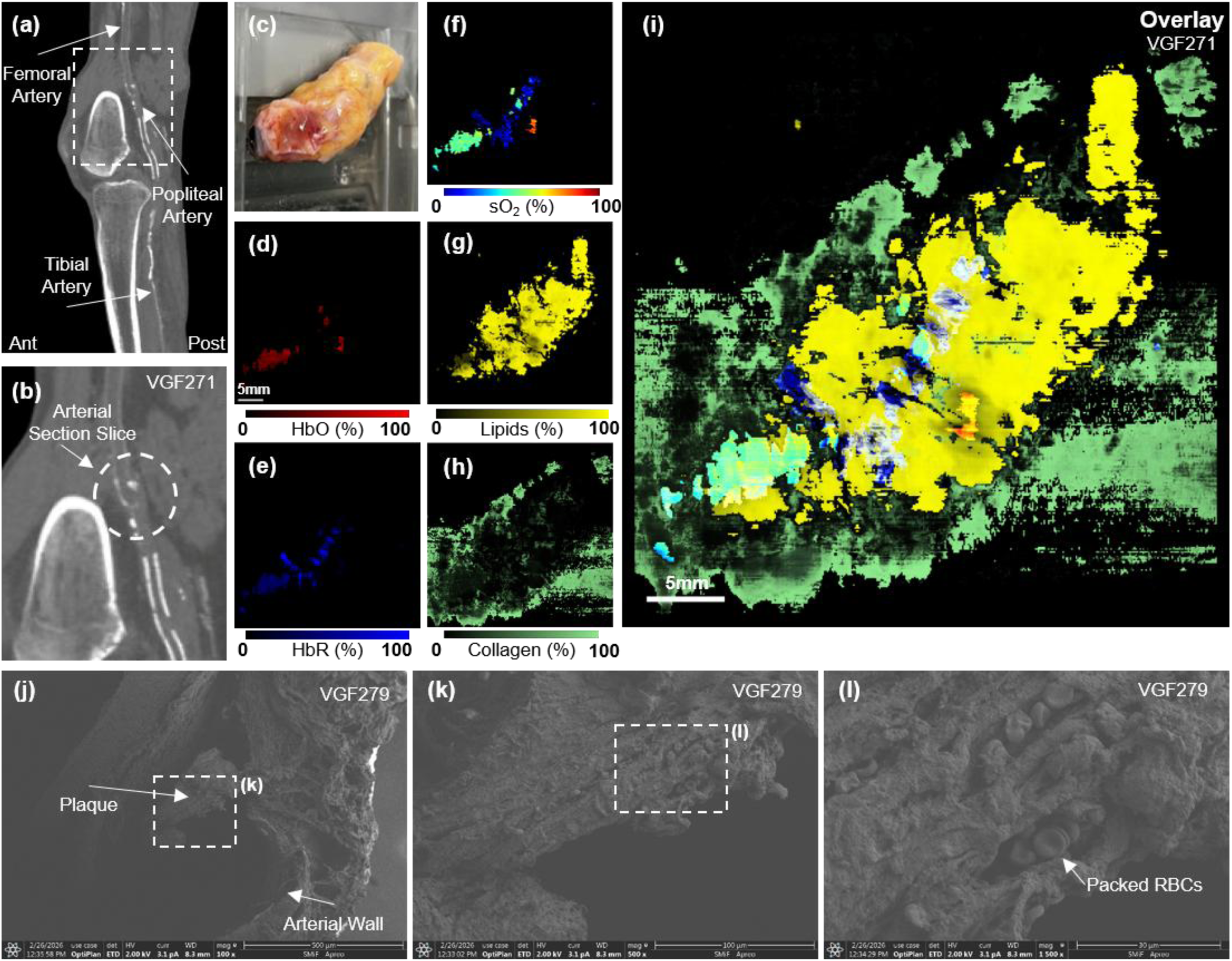
Sagittal CT images of the lower extremity (sample VGF271) showing arterial calcification and anatomical structure but lacking molecular specificity. **(a-b)** The region corresponding to the excised arterial segment is indicated. **(c)** Photograph of sample and excised artery. **(d–h)** PAF molecular maps of HbO, HbR, sO₂, lipids and collagen, revealing heterogeneous biochemical composition within the arterial segment. **(i)** Composite overlay of sO₂, lipids and collagen. PAF identifies elevated lipid signal adjacent to a localized low-sO₂ region consistent with a packed red blood cell core, indicative of thrombus and lipid-rich plaque morphology. PAF provides molecular information inaccessible to conventional CT imaging. **(j–l)** a comparable arterial specimen (sample VGF279) demonstrating plaque structure, arterial wall morphology, and densely packed red blood cells. Although not co-registered, these features support the biochemical patterns observed with PAF.

The CT scan of the patient’s lower extremity clearly indicates calcification, confirming the diagnosis of PAD. However, the lack of molecular specificity makes it difficult to identify areas of plaque buildup or potential thrombi. By contrast, PAF revealed lipid content elevated along the arterial wall, consistent with recorded atherosclerotic plaque accumulation [45]. In addition, the sO₂ map highlights a dense, oxygen-depleted region corresponding to a RBC-packed core, characteristic of thrombus composition and delineates the arterial track [46]. Together, these features provide biochemical evidence consistent with thrombus and plaque composition, information inaccessible to conventional CT imaging. The SEM imaging of comparison PAD patient’s segmented artery revealed similar high-resolution properties. However, SEM imaging requires the samples to be *ex vivo* and time-consuming steps of fixing, dehydration, and typically gold coat sputtering in order to reveal these characteristics. While not used *in vivo* in this study, PAF can already achieve similar chemical analysis *ex vivo* without the time consuming and expensive procedures of SEM.

To further evaluate the model’s robustness, we imaged another arterial specimen from a separate PAD patient. A 3D CT angiogram was available for the below-knee amputation (Figure 7a), and PAF revealed highly detailed composition of the thrombus, resolving the clot composition and plaque buildup (Figure 7b-g). These results demonstrate PAF’s capability to quantitatively and spatially resolve multiple chromophores within complex tissue.

**Figure 7:**
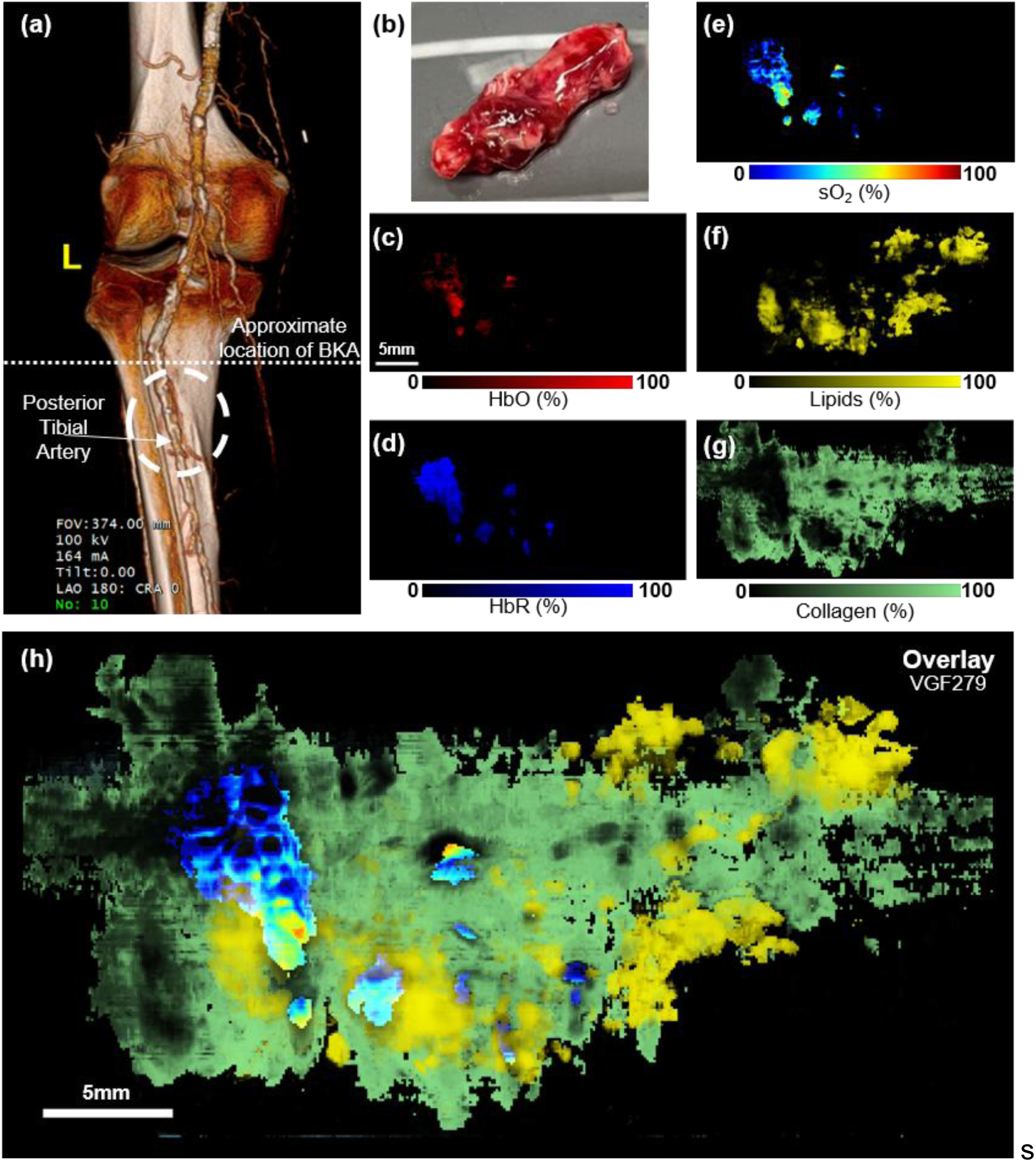
Reproducible chemical mapping of PAD using PAF. **(a)** 3D CT angiogram of the lower extremity (sample VGF279) indicating the approximate location of the excised arterial segment. While 3D CT angiogram provides high-resolution anatomical visualization of vascular structure, it lacks information on molecular composition. **(b)** Photograph of excised artery. **(c–g)** PAF maps for HbO, HbR, sO₂, lipids and collagen, revealing heterogeneous biochemical organization within the specimen. **(h)** Composite overlay highlighting lipid-rich regions and oxygen-depleted core of the thrombus.

The second sample reproduced the biochemical patterns observed in the first specimen, demonstrating that PAF enables consistent, label-free characterization of vascular disease. Notably, the two arterial specimens exhibited distinct gross appearances. The first sample showed pronounced yellow coloration along the arterial wall, consistent with lipid-rich plaque, whereas the second specimen appeared less lipid-laden. This color difference is reflected in the PAF reconstructions, which reveal substantially higher lipid signal in the yellowed regions of the first sample. By identifying expected lipid rich and collagen regions as well as sO_2_ within the same dataset, PAF provides label-free biochemical insights into plaque morphology and thrombus organization, which are key markers for assessing vascular integrity and disease progression. Together, these findings suggest that PAF can bridge the gap between anatomical imaging and molecular characterization in vascular disease.

### 5.5 Limitations and Next Steps

While this study focuses on establishing PAF as a robust *ex vivo* chemical imaging framework, several challenges must be addressed to enable reliable *in vivo* deployment.

#### 5.5.1 Training Time and Model Complexity

Although PAF inference is fast, training the RNN across large synthetic datasets remains computationally expensive (∼12 hours on an RTX 4060 Ti). Future work will explore model-compression strategies, architectural simplification, and vectorized image processing to reduce training cost and further accelerate reconstruction.

#### 5.5.2 Library Completeness

Expanding the spectral library to include more clinically relevant endogenous and exogenous absorbers will be essential for translating PAF to more complex tissues. Furthermore, because the model is trained on idealized synthetic spectra with Gaussian noise, a domain gap exists between training data and real tissues, which exhibit nonlinear fluence, wavelength-dependent attenuation, and heterogeneous molecular composition. Incorporating experimentally measured spectral fingerprints will be important for improving generalization.

#### 5.5.3 Fluence Modeling and Domain Gap

In training, we used a parametric exponential fluence prior to synthesize wavelength-dependent multiplicative distortions and to probe the network’s ability to learn non-linear mappings from distorted fingerprints to concentration vectors. While this prior affords controlled experiments, real tissue light transport is governed by complex absorption and scattering (and therefore multiple scattering, angular redistribution, and geometry) and is not fully captured by 1D exponential attenuation. Recent work has demonstrated that absorbing molecules such as indocyanine green (ICG) can act as optical clearing agents *in vivo,* improving penetration depth and contrast by modulating tissue optical properties, suggesting a promising complementary strategy for mitigating fluence related distortions in deep tissue photoacoustic imaging [47]. Still, a domain gap exists between the synthetic training distribution and *in vivo* fluence distribution. Mitigation strategies include: (i) augmenting training with diffusion-approximation or Monte Carlo fluence models, (ii) fine-tuning the network on a small set of experimentally measured fingerprints, or (iii) integrating fluence-estimation modules into the reconstruction pipeline. These paths are tractable and represent immediate next steps for *in vivo* translation. Future work will focus on integrating experimentally measured fingerprints and physics-informed simulations to further close this domain gap.

#### 5.5.4 Depth, Fluence and Noise in Complex Media

The multicomponent phantom experiments performed in this work had limited depths (<1cm) and do not capture tissue scattering, acoustic attenuation or physiological motion present *in vivo*. Validation in deeper tissue-mimicking phantoms and small-animal *in vivo* studies will be necessary to evaluate performance under realistic biological conditions.

The *ex vivo* mouse liver experiment had a limited sample size, providing only qualitative demonstrations rather than statistically meaningful biomarkers. While the *ex vivo* human artery experiment in this study provide an important step toward clinical relevance, the arteries do not replicate *in vivo* optical, hemodynamic, or physiological conditions. In particular, the absence of blood perfusion and rapid deoxygenation of static blood alter hemoglobin oxygenation compared to living systems. Nevertheless, larger sO₂ differences *in vivo* may actually enhance contrast between thrombus and surrounding blood. Future studies will incorporate larger sample sizes, *in vivo* imaging, and deeper tissue models to evaluate PAF under physiologically realistic conditions. Recent studies combining absorbing dyes with photoacoustic microscopy have demonstrated improved imaging depth and sensitivity *in vivo,* highlighting the potential for integrating physical contrast enhancement strategies with data-driven unmixing frameworks such as PAF for next-generation molecular imaging [48]. By systematically addressing these limitations and next steps, we anticipate that PAF will mature into a reliable and generalizable tool for quantitative deep-tissue molecular imaging.

## 6. Conclusions and Future Work

In this study, we introduced photoacoustic fingerprinting (PAF), a novel machine learning approach for deep-tissue molecular unmixing, designed to overcome the limitations of conventional linear reconstruction techniques. Built upon a RNN framework, PAF learns complex wavelength-dependent variations from a unique “photoacoustic fingerprint,” enabling robust and noise-tolerant spectral unmixing.

Through comprehensive synthetic data analysis, PAF demonstrated superior accuracy in reconstructing molecular concentrations across SNR and fluence variations, outperforming NNLS unmixing. Specifically, PAF improved R^2^, reduced RMSE, and exhibited a fitted slope closer to unity compared to NNLS. The AAE decreased across all molecules, with the largest improvement in collagen. Phantom validation of chromophore groups confirmed increasing sampled wavelengths increased NNLS accuracy. The overdetermined set confirmed PAF’s ability in a low SNR environment to reconstruct concentrations more accurate than NNLS.

Beyond synthetic and phantom validations, we further demonstrated PAF’s capability in biologically and clinically relevant tissues. In *ex vivo* mouse liver analysis, PAF quantitatively detected lipid accumulation associated with HFD–induced steatosis. This result underscores PAF’s sensitivity to biochemical variations arising from metabolic alteration and establishes its potential for assessing endogenous composition without exogenous contrast agents.

Extending this framework to human arterial samples, PAF successfully identified distinct molecular signatures of thrombus and plaque formation in patients with PAD. Elevated lipid and deoxygenated hemoglobin signals delineated the thrombus core. The successful identification of lipid-rich plaque and thrombus signatures in human PAD arteries further highlights the clinical relevance of PAF for vascular disease assessment. SEM imaging of a comparable sample provided high-resolution confirmation of plaque and RBC-packed core. To our knowledge, this work represents the first application of a RNN–based spectral fingerprinting approach to *ex vivo* human vascular tissue.

Together, these results highlight PAF’s ability to quantitatively resolve endogenous molecular distributions across diverse biological systems, bridging the gap between controlled experimental validation and clinically meaningful tissue analysis. As molecular-resolution approaches continue to emerge, PAF may complement anatomical imaging modalities by providing biochemical context that is otherwise inaccessible. More broadly, PAF introduces a generalizable paradigm for photoacoustic chemical imaging. Molecules with known absorption spectra, in principle, can be incorporated into the fingerprint library, enabling targeted or exploratory chemical analysis without redesigning the reconstruction framework. This flexibility positions PAF as a foundation for future chemically specific photoacoustic imaging studies across *ex vivo*, preclinical, and ultimately *in vivo* applications.

Future work will focus on addressing current limitations to facilitate clinical translation. This includes exploring model-compression techniques to reduce the substantial training cost of the RNN while also investigating new ways to generate synthetic samples with fluence variations. As well as this, expanding the spectral library to include a broader range of endogenous and exogenous agents is crucial to enhance the model’s applicability to *in vivo* tissues. Finally, comprehensive validation in deeper, tissue-mimicking phantoms and small animal *in vivo* experiments will be essential to assess PAF’s performance under more realistic conditions, including optical attenuation, acoustic attenuation and motion. Collectively, we expect that PAF will evolve into a powerful and generalizable framework for quantitative, label-free molecular imaging with clear potential for clinical translation.

## 7. Declarations

### 7.1 Funding

This work was partially sponsored by the United States National Institutes of Health (NIH) grants R01EB037095, RF1 NS115581, R01 NS111039, R01 EB028143, R01 DK139109, R01 DK052985, R01 MH135932; The United States National Science Foundation (NSF) CAREER award 2144788; American Heart Association Collaborative Science Award (25CSA1417550); Duke Gilhuly Acceleration Grant; Duke University Pratt Beyond the Horizon Grant; Eli Lilly Research Award Program; Chan Zuckerberg Initiative Grant (2024-349531); Duke University DST Spark Seed Grant; Duke Coulter Translational Grant; North Carolina Biotechnology Center Triangle Research Grant (2024-TRG-0041).

### 7.2 Competing interests

J.Y. and L.M. have financial interests with Lumius Imaging, Inc., which did not support this work. Other authors declare no competing interests.

### 7.3 Author Contributions

C.M. and J.Y. conceived the idea. C.M. designed and trained the PAF-RNN. L.M. designed and built the DAT imaging system. C.M. performed all imaging studies. C.M. and D.W. performed multi-component phantom imaging and analysis. C.M. and L.M. performed *ex vivo* liver imaging. L.M. contributed to liver data analysis. S.F. sourced and monitored the human artery imaging study. C.M. performed artery imaging. A.Y., V.T.N., S.W.C., L.M., D.W., S.F. and C.M. contributed to data analysis. C.M. and J.Y. wrote the manuscript with input from all authors. J.Y. supervised the whole study. All authors reviewed and approved the final manuscript.

### 7.4 Data Availability

The code and data supporting the findings of this study are available from the corresponding author upon reasonable request.

## 8. Ethics Approval and Consent to Participate

All animal experiments were conducted in compliance with Institutional Animal Care and Use Committee (IACUC) guidelines at Duke University (Duke IACUC protocol: 60681). Human arterial samples used in this manuscript were collected using an institutional review board (IRB) approved protocol (Duke IRB#: Pro00065709).

## Acknowledgements

The authors thank Dr. Gowthami Arepally, Dr. Shrike Zhang, Joseph Yang, Shruthi Srinivasan, Wren Wightman and Max Golovsky for valuable discussions related to this work.

## Notes

### Competing Interest Statement

The authors have declared no competing interest.

